# scIDPMs: single-cell RNA-seq imputation using diffusion probabilistic models

**DOI:** 10.1101/2024.02.29.582870

**Authors:** Zhiqiang Zhang, Lin Liu

## Abstract

Single-cell RNA sequencing (scRNA-seq) technology is a high-throughput sequencing analysis method that enables the sequencing of mRNA in individual cells, thereby facilitating a more precise understanding of cellular gene expression and metabolic products. This approach reveals cell function and characteristics, making it widely applicable in biological research. However, scRNA-seq data often suffers from false zero values known as dropout events due to limitations in sequencing technology. These dropout events not only mask true gene expression levels but also significantly impact downstream analysis accuracy and reliability. To address this challenge, numerous computational approaches have been proposed for imputing missing gene expression values. Nevertheless, existing imputation methods struggle to fully capture the distribution of dropout values due to the high sparsity of scRNA-seq data and the complexity and randomness associated with gene expression patterns. Recently, probabilistic diffusion models have emerged as deep generative models capable of accurately restoring probability density distributions in domains such as image and audio processing. In this paper, we propose a method called scIDPMs, which utilizes conditional diffusion probabilistic models to impute scRNA-seq data. scIDPMs first identifies dropout sites based on the characteristics of cellular gene expression and then infers the dropout values by conditioning on the available gene expression values, which provide context information for the dropout values. To effectively capture the global features of gene expression profiles, scIDPMs employs a deep neural network with an attention mechanism to optimize the objective function. The performance of scIDPMs was evaluated using both simulated and real scRNA-seq datasets, and compared with eight other imputation methods. The experimental results clearly demonstrated that, in comparison to alternative approaches, scIDPMs exhibited exceptional performance in recovering biologically meaningful gene expression values and enhancing various downstream analyses.

## 1 INTRODUCTION

As the fundamental unit of life, cells exhibit heterogeneity and complex interactions within biological systems. The diversity of gene expression is a key factor contributing to the variation in cell morphology and function. While bulk RNA-seq technology provides information on average gene expression levels across entire cellular subpopulations, it fails to capture cellular heterogeneity. In contrast, single-cell RNA sequencing (scRNA-seq) enables construction of gene expression profiles for individual cells, providing a more precise understanding of gene expression at the single-cell level. This technology presents new opportunities for exploring heterogeneity and functional differences within cell populations and has been widely applied in fields such as cancer research, cell development, and immunology [1]. However, scRNA-seq data contain numerous zero values due to both biological reasons (i.e., genes not expressed or expressed at low levels) and limitations in sequencing technology (e.g., cDNA amplification deviation or insufficient sequencing depth), resulting in dropout events [2, 3]. These events pose significant challenges for downstream analysis tasks such as identifying intercellular gene expression differences or characterizing cell types [4].

To tackle the issue of missing gene expression values resulting from dropout events, a range of methodologies have been proposed, encompassing model-based and deep learning-based methods. Model-based methods utilize prior knowledge such as existing gene expression values, cell relationships, and gene relationships to establish probabilistic models for estimating missing values and uncovering gene expression patterns in single-cell transcriptomic data. For instance, kNN-smoothing [5] updates the expression values of each cell by averaging the expression information from its *K* nearest neighboring cells. MAGIC [6] constructs a similarity graph of cells and propagates gene expression information between similar cells to infer missing values. SAVER [7] models sequencing data using a probabilistic approach and employs Bayesian statistical inference for recovery purposes. Although these methods have achieved some success in imputing scRNA-seq data, they tend to globally adjust the expression levels of all genes, potentially distorting true gene expressions and introducing noise in downstream analysis - leading to an over-smooth problem where imputed data perform worse than unimputed data. To tackle this issue, scImpute [8] first identifies dropout sites and then imputes dropout values based on gene expression information from similar cells; however, it may also reduce random differences in gene expressions between cells [9].

With the robust fitting capability of deep neural networks, an increasing number of deep learning-based imputation methods for scRNA-seq data have been proposed. These methods primarily rely on specific neural network structures, such as convolutional neural networks (CNN) [10] and graph neural networks (GNN) [11], to extract data features and restore data distribution from latent space using generative models like AutoEncoders (AEs) [12], Variational Auto-Encoders (VAEs) [13], and Generative Adversarial Networks (GANs) [14]. For instance, AutoImpute [15] employs VAEs to restore the initial distribution of sparse matrices, while scIGANs [16] reshape individual cell expression profiles into images by treating missing gene expression values as missing pixels and use GANs to recover the expression data. However, this method faces limitations due to the limited receptive field of convolutional operations, making it challenging to capture global data features. To address this issue, GNNImpute [17] utilizes graph attention network (GAT) to extract cell features and employs AEs for reconstructing the expression matrix. SCDD [18] combines graph convolutional network (GCN) with AEs to reduce noise introduced during imputation in order to alleviate over-smoothing problems. LSH-GAN [19] applies locality sensitive hashing for sampling processed expression data and then trains GANs using the sampled data to generate new samples, thereby addressing issues related to fewer cell samples in downstream analysis tasks. Additionally, scGCC [20] leverages GAT for learning low-dimensional representations of the data while scGCL [21] combines graph contrastive learning with AEs for imputing missing values. scGGAN [22] is a GANs model based on GCN. It initially constructs a gene relation network using additional genomic sequencing data and subsequently utilizes this network, along with the expression matrix of scRNA-seq, as training data to learn and restore the underlying distribution of the dataset. Despite the evaluation conducted by Dai et al. [23], which demonstrates superior performance of deep learning-based imputation methods over model-based imputation approaches, there still exists potential for further enhancement.

Typically, scRNA-seq data contains gene expression information for thousands of genes across a large number of cells, resulting in high dimensionality [24]. Moreover, scRNA-seq data is characterized by high sparsity and noise levels that are influenced by various factors such as sequencing technology, environment, and physiological states. These features present challenges for model-based imputation methods to accurately capture the underlying data distribution. Recently, diffusion probabilistic models (DPMs) [25, 26], as a class of likelihood-based generative models, have emerged as the new state-of-the-art family of deep generative models [27]. DPMs are capable of accurately restoring the ground-truth distribution of target data on hidden variables with dimensions identical to those of the original data while maintaining diversity in generated samples. Compared to other generative models such as VAEs and GANs, DPMs estimate target distributions based on variational inference with a solid theoretical foundation, flexible model structure, and stable training process. As a result, they achieve excellent performance in computer vision, natural language processing, multimodal learning and other fields [28]. The existing deep learning-based methods employ various deep neural network models, such as CNN and GNN, to partially extract local features from gene expression sequences; however, acquiring comprehensive global features remains challenging. Simultaneously, advanced language models built upon the Transformer [29] architecture, such as ChatGPT [30] and LaMDA [31], have made remarkable strides in the field of natural language processing. These models leverage deep neural networks that incorporate attention mechanisms to effectively capture long-range dependencies within sequences. Inspired by Zheng et al. [32], who employed DPMs for the purpose of imputing missing values in tabular data, this paper proposes scIDPMs, a framework based on conditional DPMs designed specifically for scRNA-seq imputation (Figure 1). ScIDPMs constructs a conditional deep neural network model with attention mechanism based on the powerful inference algorithm of DPMs to restore the distribution of complex target data, utilizing existing gene expression values as conditions for inferring dropout value distributions. Specifically, scIDPMs identifies dropout sites by referencing gene expression patterns in similar cells and employs a self-supervised training method to divide existing gene expression values into conditional observation values and imputation targets. These two sets are then used as training data for optimizing the parameters of the conditional DPMs model with attention mechanism. Finally, the trained conditional DPMs model takes existing gene expression values as conditional observations and gradually restores the ground-truth distribution of dropout values through an inference algorithm based on DPMs.

**Figure 1.**
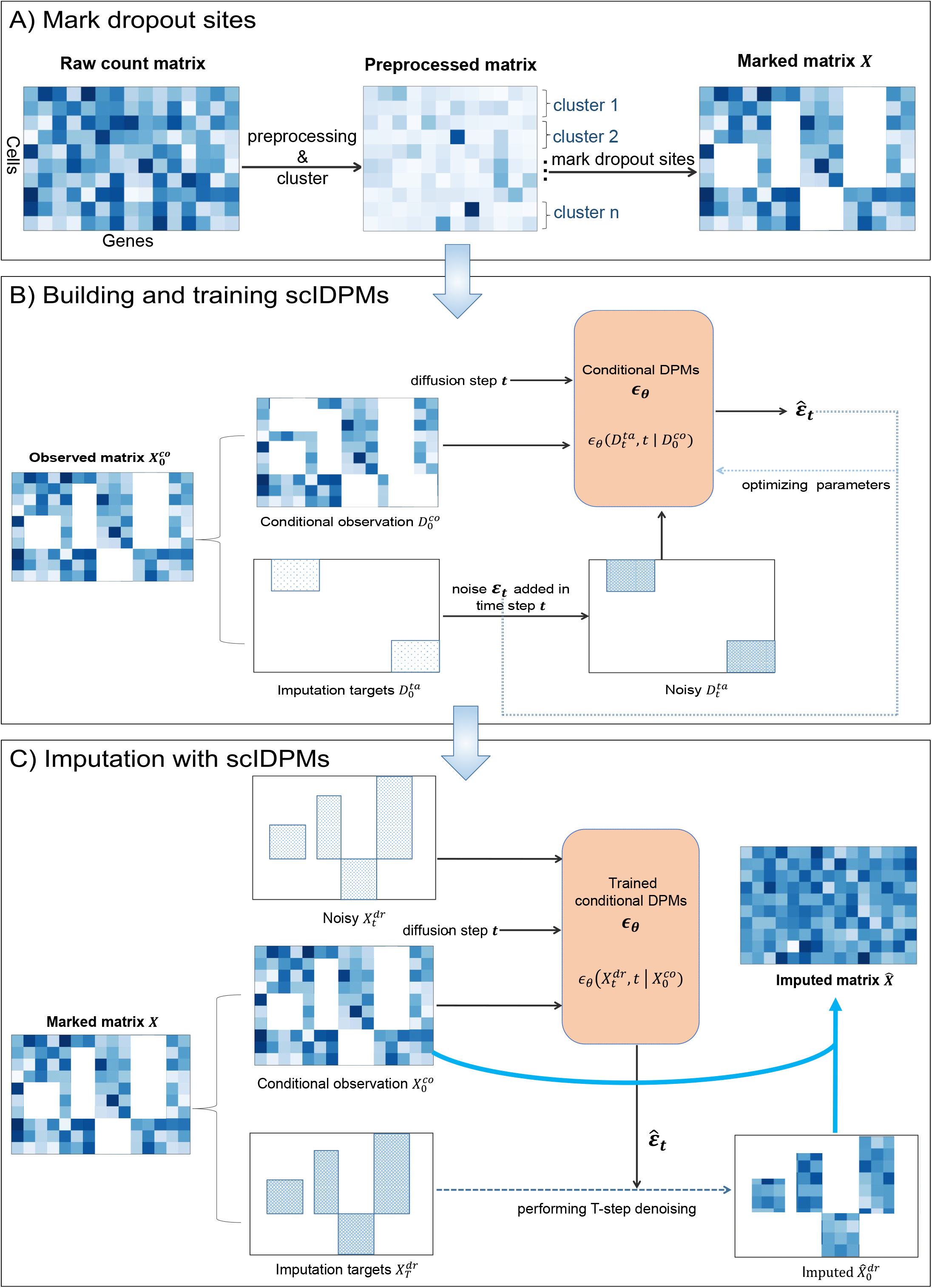
Overview of scIDPMs for scRNA-seq data imputation. (**A**)Preprocessing the raw data and identifying dropout sites based on intercellular relationships. (**B**)Constructing training datasets for conditional DPMs using existing gene expression values. (**C**)Imputing dropout values by utilizing the trained conditional DPMs with existing gene expression values as conditional inputs.

Compared to existing imputation methods, scIDPMs possess several advantages. Firstly, leveraging the inherent strengths of DPMs inference algorithm, scIDPMs does not rely on a specific statistical model for the distribution of target data. This allows it to better fit the distribution of dropout values. Secondly, scIDPMs exclusively imputes expression values affected by dropout events without altering the original gene expression values and biological zeros. Consequently, it avoids introducing additional noise in downstream analysis. Thirdly, scIDPMs employs a deep neural network model with an attention mechanism to capture comprehensive context information from single-cell gene expression profiles. This enables it to extract features more comprehensively from gene expression sequences. Lastly, unlike other methods that require supplementary information such as corresponding bulk RNA-seq data or genome sequence data for imputation purposes, scIDPMs can complete interpolation independently. In this study, we systematically evaluate the performance of scIDPMs using simulated and real scRNA-seq datasets and compare it with eight other imputation methods. Our experimental results demonstrate that scIDPMs exhibit significant advantages in restoring gene expression levels and improving downstream tasks including cell clustering and differential expression analysis.

## 2 Methods

### 2.1 Notation

The symbols presented in this section are as follows: the expression matrix after quality control is denoted as *X ∈* ℝ^*n×m*^, where *n* represents the number of cells and *m* represents the number of genes. The part with dropout is denoted as 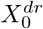, with its corresponding indices set as *M*^*dr*^; the remaining part is denoted as 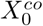, with its corresponding indices set as *M*^*co*^. The objective of this paper is to construct the model 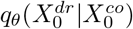 for estimating 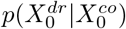 .

### 2.2 Mark dropout sites

Determining dropout values in gene expression is a critical step, and existing methods can be classified into statistical analysis-based and biology-based approaches. However, due to the inherent noise and complexity of gene expression data, accurately modeling the distribution of dropout sites using statistical probability models poses challenges. Therefore, this study proposes inferring dropout sites by leveraging similar cell types’ gene expression information from a biological perspective (Figure 1 A). Specifically, for scRNA-seq data with known cell types, cells of the same type are grouped together based on their expressions of each gene. If a gene exhibits both expressed and unexpressed cells within the group, its unexpressed location is identified as a dropout site and recorded.

For scRNA-seq data without cell type information, the clustering results of cells can be utilized as class labels for each individual cell, and the aforementioned steps are repeated to annotate dropout events. The detailed algorithm is presented in Algorithm 1 . The process of clustering scRNA-seq data can be summarized as follows: firstly, normalizing the expression values of each cell to a range between 0 and 1 and applying logarithmic transformation to mitigate discrepancies in expression levels; subsequently, employing the Uniform Manifold Approximation and Projection (UMAP) [33] algorithm to compute cellular neighborhood relationships. UMAP is a non-linear dimensionality reduction technique that effectively maps high-dimensional data onto a lower-dimensional space while preserving local structural information between cells. Lastly, comprehensive cellular cluster analysis is performed using clustering algorithms such as Leiden [34].

#### Algorithm 1

Mark dropout sites

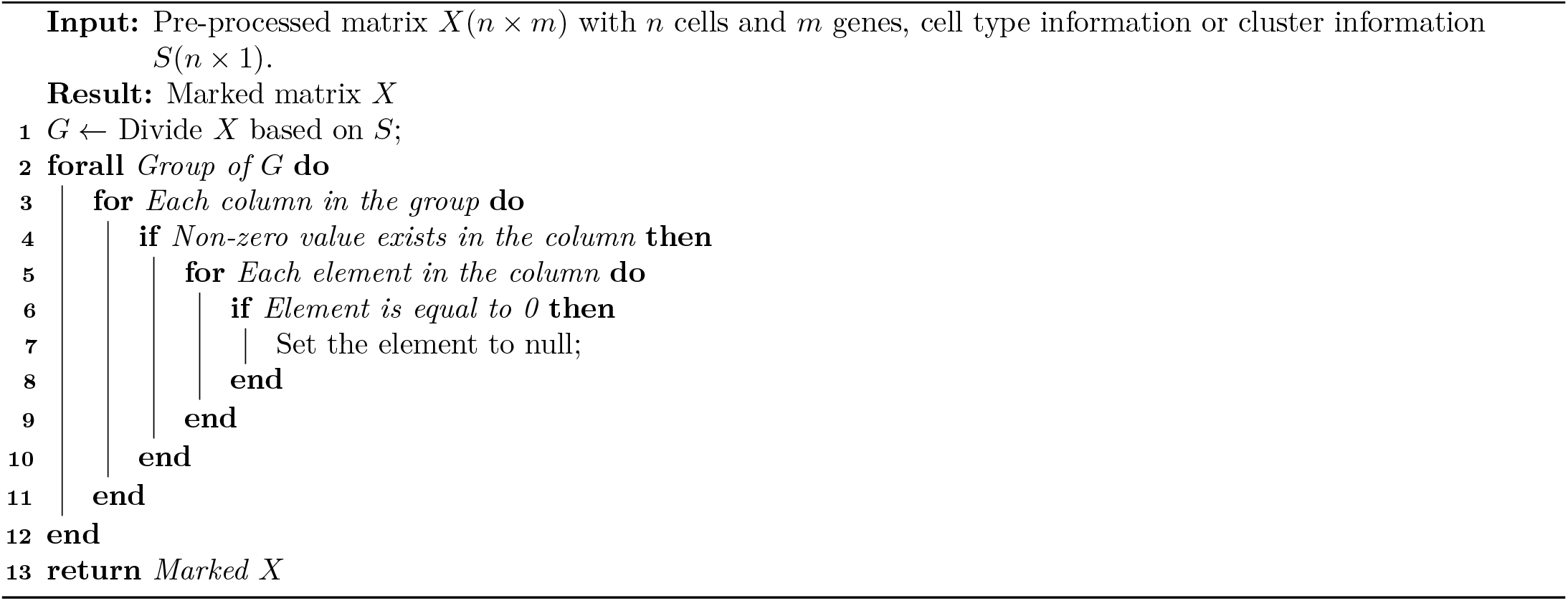

### 2.3 Building and training scIDPMs

#### 2.3.1 Diffusion probabilistic models

This section first gives a brief description of vanilla DPMs, which provides a solid theoretical foundation for scIDPMs. DPMs consists of two parameterized Markov chains, the forward process and the reverse process [35]. Assuming that the distribution of the training data is *x*_0_ *∼ q*_0_(*x*_0_), the goal of DPMs is to learn a model distribution *p*_*θ*_(*x*_0_) that approximates the marginal distribution *q*_0_(*x*_0_). Specifically, in the forward process, DPMs add a fixed increasing variance schedule *β*_1_, …, *β*_*T*_ with *β*_*t*_ *∼* (0, 1) Gaussian noise to the data at each time step *t*. After T steps of noise addition, the distribution of the data eventually tends to the standard Gaussian distribution. The data *x*_*t*_ after the *t* step of noise addition to the original data satisfies:

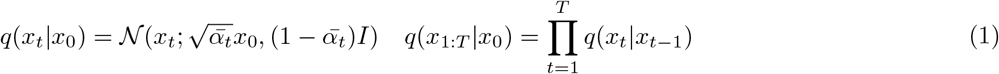

where *α*_*t*_ := 1 *− β*_*t*_, 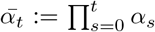, *t ∈* [0, *T*] is the number of steps for adding noise. It is worth noting that in the case of given *x*_0_, the reverse diffusion process can be inferred by using Bayes’ rule and reparameterization trick:

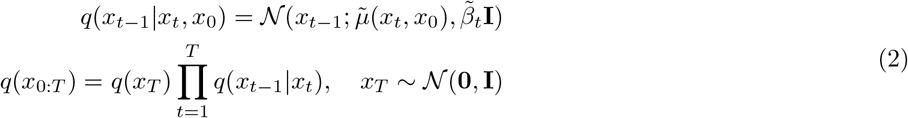

where 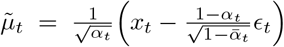, *ϵ* is the noise sampled randomly from the normal distribution at time *t*, and 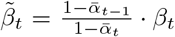 . If the exact reverse distribution *q*(*x*_*t−*1_|*x*_*t*_) is known, then the ground truth distribution *q*_0_(*x*_0_) of the original data can be inferred from *x*_*T*_ *∼ 𝒩* (0, **I**). So the modeling objective is:

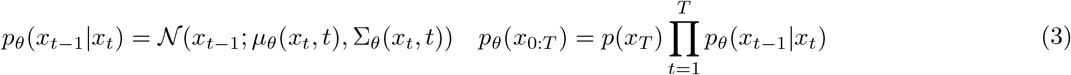

The optimization goal can be constructed by the lower bound of the variation of the negative log likelihood function of the model *p*_*θ*_(*x*_0_):

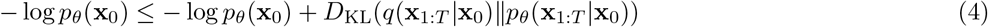

Splitting the Kullback-Leibler divergence of the two joint probability distributions in Eq.4 and disregarding the constant term, we obtain the optimization objective of the model as follows:

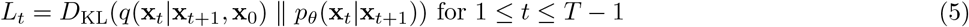

The new optimization goal is obtained by substituting the reverse distribution obtained from Eq.2:

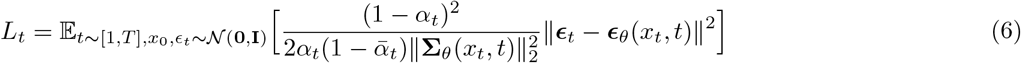

Ho et al. [26] discovered that the optimal approach is to disregard the constant weight term in Eq. 6 and solely predict the noise *ϵ*_*t*_ introduced during the *t*-th step of the forward diffusion process. Consequently, parameter *θ* is optimized by minimizing the following objectives:

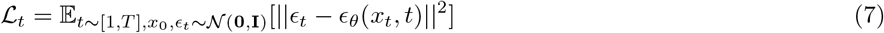

Therefore, the model *ϵ*_*θ*_ is a function with *x*_*t*_ and *t* as inputs, and its goal is to predict the Gaussian noise sampled at *t* time *ϵ*_*t*_, and update the parameters by mean square error (MSE) loss between the true noise and predicted noise. Once trained, it can randomly sample *x*_*T*_ *∼ 𝒩* (**0, I**), and iterated T steps of denoising process according to Eq. 8, and finally restore the distribution of *x*_0_.

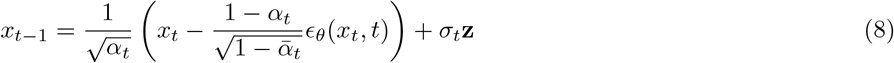

where **z** *∼ 𝒩* (**0, I**) if *t >* 1 otherwise **z** = 0.

#### 2.3.2 Training algorithm and structure of scIDPMs

To restore the distribution of 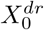 based on DPMs inference algorithm, the optimal approach is to construct conditional DPMs 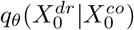 to estimate 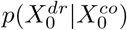, thereby enabling the inference of 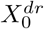 through contextual information from the entire gene expression sequence. However, due to the unknown ground-truth dropout values, this study employs a self-supervised training method that segregates 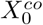 into conditional observation values and imputation targets (Figure 1 B). The specific division method involves randomly selecting a certain proportion of values as imputation targets, denoted as 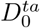, with corresponding indices *m*^*ta*^ ; while considering the remaining values as conditional observation values, denoted as 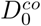, with corresponding indices *m*^*co*^. According to Eq. 1, the noisy 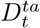 satisfies:

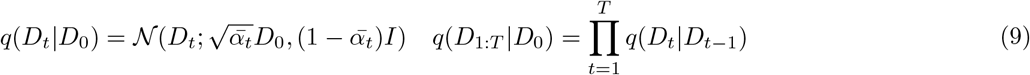

The modeling objectives of the aforementioned conditional DPMs are formulated by extending Eq. 3:

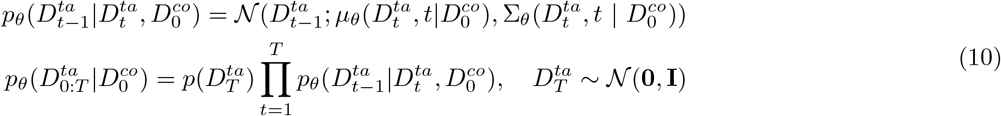

Since the model solely aims to estimate the noise *ϵ*_*t*_ added to 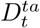 at time *t*, the optimization objective of the model in this stage is determined based on previously recorded indices information:

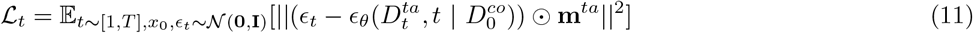

##### Algorithm 2

Training of scIDPMs

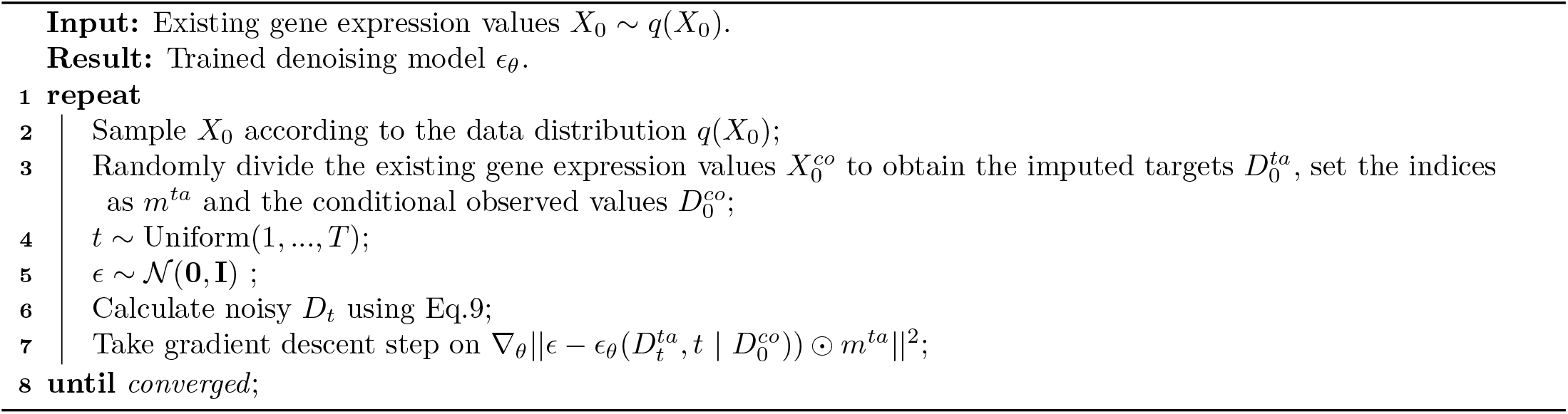

The training procedure of scIDPMs is delineated in Algorithm 2.

The key to effectively addressing optimization objectives in deep learning lies in constructing appropriate network architectures for extracting data features, thereby facilitating enhanced model parameter optimization. The deep neural network architecture utilized by scIDPMs is illustrated in Figure 2. The training data for the model comprises three components: noise is added to 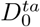 through Eq. 1 to obtain noisy 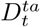, which is then concatenated with normalized 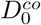 as the ‘feature’ part of the training data; to better extract contextual information from individual cell expression values, the indicator matrix *m*^*co*^, *m*^*ta*^, and coordinate information are combined as ‘conditional information’ and input into the model; finally, in order to capture the relationship between noisy 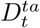 and the noise *ϵ*_*t*_ added in step *t* at different time steps, use the following 128-dimensional embedding following previous works [36] and then input it into the model.

**Figure 2.**
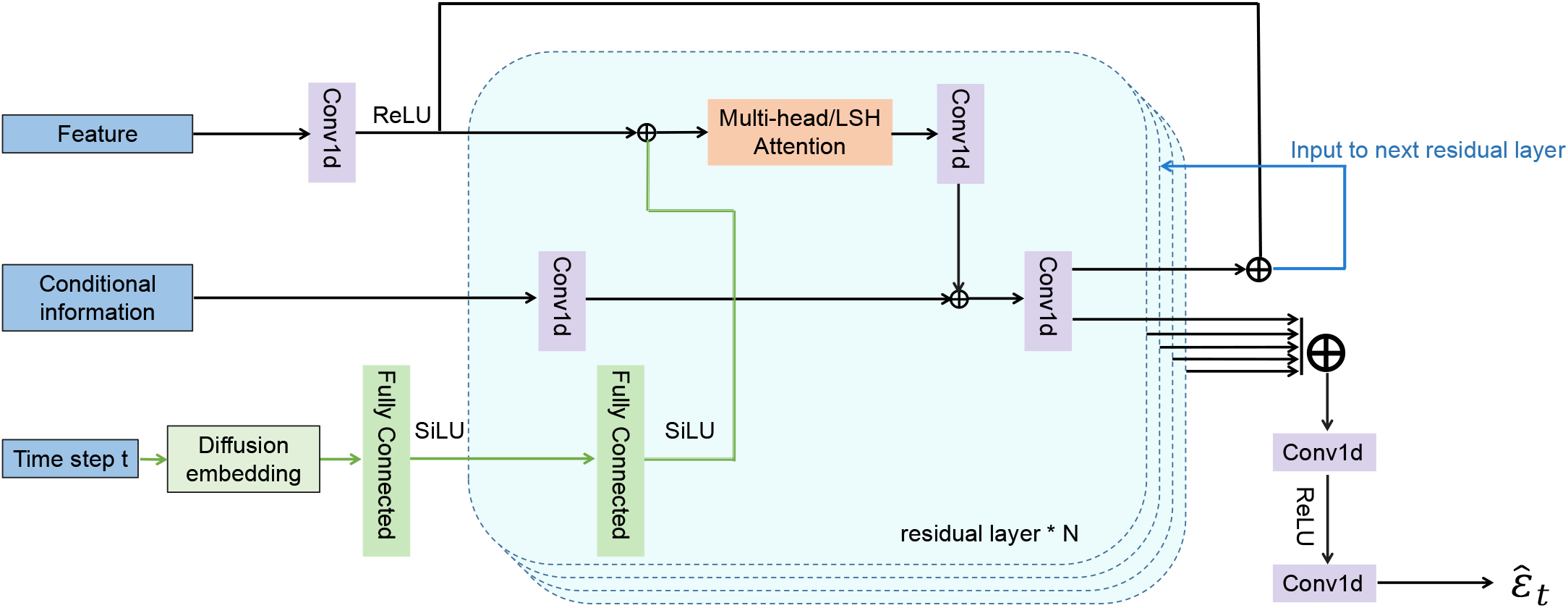
The deep neural network architecture employed by scIDPMs.

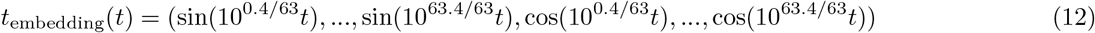

#### 2.3.3 Attention mechanism in scIDPMs

Unlike previous deep neural network-based interpolation methods, scIDPMs extract data features by constructing residual blocks [37] that contain Conv1 *×* 1 [38] and attention mechanisms [39]. Conv1 *×* 1 is a one-dimensional convolutional neural network that uses convolution kernels to perform operations on expressed data through sliding window mechanism, thereby extracting local features. Multiple convolution layers can enable the model to learn more complex feature combinations, improving its expressiveness and generalization ability. Additionally, Conv1 *×* 1 ensures that the data feature map remains unchanged by adjusting the kernel size to expand the data channel and enhance interaction between channels.

Compared to other models for sequence processing, the Transformer effectively captures contextual dependencies in a sequence by assigning distinct attention weights to its elements, thereby enhancing its ability to capture global information. The multi-head self-attention mechanism plays a pivotal role in this model. For a sequence containing *n* elements, each element is mapped to query (*Q*), key (*K*), and value (*V*) through three learnable parameter matrices, *W*^*Q*^, *W*^*K*^, and *W*^*V*^, all of which have a dimension of *d*_*k*_. The attention weights for each element, also known as scaled dot-product attention, can be obtained using Eq. 13 .

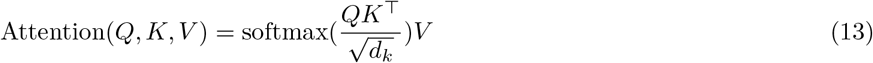

Instead of computing attention only once, the multi-head mechanism concurrently performs multiple iterations of scaled dot-product attention. The resulting attention outputs from each head are then concatenated and linearly transformed to achieve the desired dimensions (Eq. 14).

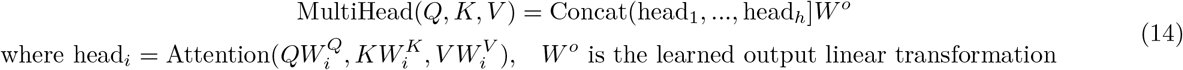

However, for input sequences of considerable length, the computation of multi-head self-attention consumes significant memory space and computational resources. Moreover, a substantial portion of scalar scores between *Q* and *K* are either zero or close to zero, rendering it unnecessary to compute attention scores for all positions. To address this issue, Reformer [40] incorporates locality sensitive hashing (LSH) into the attention mechanism by employing a multi-plane LSH scheme to identify the nearest neighbor vector *k*_*j*_ *∈ K* that is closest to *q*_*i*_ *∈ Q*. By performing multiple random mappings, *Q* and *K* are partitioned into distinct buckets where only attention weights within the same bucket are calculated. Consequently, this approach reduces complexity from *O*(*n*^2^) to *O*(*n* log *n*). Nevertheless, it should be noted that the LSH attention mechanism employs hash functions for approximate calculation which may introduce certain errors.

In the task of imputing scRNA-seq data, the gene expression information of individual cells continues to increase, resulting in a continuous expansion of the gene expression scale. To address this challenge, we constructed deep neural network models utilizing both multi-head attention mechanism and LSH attention mechanism, and compared their performance on imputation results. Our aim is to identify a more suitable imputation method for handling large-scale single-cell transcriptomic data. Specifically, we implemented the multi-head attention mechanism using Transformer encoder architecture comprising a multi-head attention layer, fully-connected layers, and layer normalization. The LSH attention mechanism was implemented by stacking LSH attention layers. Additionally, scIDPMs adopted residual connections to mitigate gradient vanishing during training and enhance model expressiveness by combining original data features with those extracted by deep networks.

### 2.4 Imputation with scIDPMs

After the completion of model training, the normalized existing gene expression value 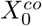 is utilized as conditional inputs to infer missing values through the trained conditional DPMs (Figure 1 C).

Specifically, we initialize the denoising object by randomly sampling 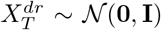, and then subject to *t* steps of noise addition applied to 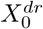, resulting in 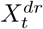 along with its index information *M*^*dr*^. Additionally, the diffusion step *t* and conditional information 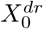 along with its index *M*^*co*^ are provided as inputs to the trained model. The model outputs 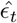 representing the noise added in the *t*-th step. The denoising process is iteratively performed for T steps according to the following procedure:

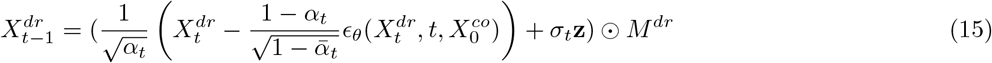

where **z** *∼ 𝒩* (**0, I**) if *t >* 1,otherwise **z** = 0. To mitigate the impact of random factors, we conducted 200 denoising inferences for each sample by employing single-cell gene expression profiles. The predicted value of the model was obtained by averaging all results. The final interpolation result *hatX* was derived from merging the denormalized 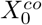 with the denoised 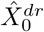 matrix. Algorithm 3 outlines the imputation process of scIDPMs.

#### Algorithm 3

Imputation with scIDPMs

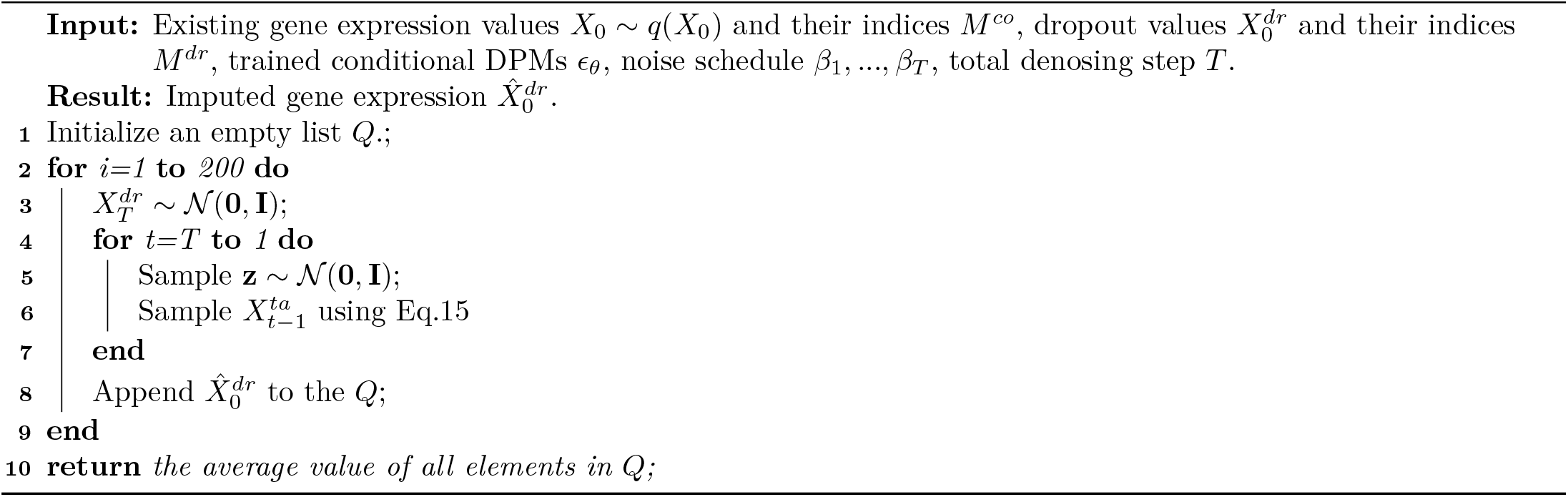

### 2.5 Experimental settings

The PyTorch [41] machine learning framework was employed in this study to construct and train the model, while the Python package Scanpy [42] was utilized for preprocessing and downstream analysis of single-cell RNA sequencing data. Unless otherwise specified, default hyperparameters were adopted during model training: 10% randomly masked existing gene expression values were used as imputation targets when constructing the training data; a cosine variance schedule, as described in a previous study [43], was applied to smoothly introduce noise into the data through forward noise process; T = 300 forward noise steps were performed; a batch size of 32 was used; the number of training epochs is 500; an initial learning rate of 0.0005 was set and dynamically adjusted by a custom learning rate scheduler throughout the training process; finally, the residual block layer is set to 4.

In order to comprehensively investigate the performance of scIDPMs, this study conducted a comparative analysis with eight state-of-the-art scRNA-seq imputation methods, namely kNN-smoothing, MAGIC, SAVER, scImpute, scIGANs, GNNImpute, SCDD, and scGCC. These methods were implemented using their default parameters as provided in the shared code for imputing scRNA-seq data without any specific instructions. All experiments were performed on an Ubuntu server equipped with an NVIDIA RTX 3080Ti GPU and 32 GB memory.

## 3 RESULTS

### 3.1 scIDPMs effectively recovers gene expression

An ideal imputation method should accurately restore dropout values. However, due to the unknown real dropout sites and values, Hou et al. [44] compared the performance of interpolation methods in restoring biological expression by calculating the correlation between bulk RNA-seq data of the same cell line and scRNA-seq data before and after imputation. Nevertheless, many scRNA-seq datasets lack corresponding bulk RNA-seq data as a reference, and significant differences exist in gene expression levels between these two types of sequencing due to their distinct principles. Considering these factors, this study evaluates the performance of scIDPMs in restoring missing gene expression values using a simulated dataset disclosed by Dai et al. [23] . The simulated dataset is generated using R package Splatter [45] for simulating scRNA-seq data. By adjusting the parameters of `dropout.mid`, a counting matrix containing four types of cells with an equal number in each type (500(cells) *×* 1000(genes)) is constructed into six datasets with varying `dropout␣rates` (Table 1 provides specific parameters for each dataset). Since the simulated dataset includes real dropout sites and gene expression matrices, this paper assesses the performance of scIDPMs in restoring gene expression values through quantification and visualization approaches while comparing them with other methods.

**Table 1.** The synopsis of the simulated scRNA-seq datasets.

| Dataset name | Dropout Rate | Zero Rate |
| --- | --- | --- |
| simulated dataset 1 | 0.55 | 0.78 |
| simulated dataset 2 | 0.47 | 0.71 |
| simulated dataset 3 | 0.38 | 0.63 |
| simulated dataset 4 | 0.30 | 0.55 |
| simulated dataset 5 | 0.23 | 0.48 |
| simulated dataset 6 | 0.16 | 0.42 |

For the purpose of quantification, we employ Root Mean Square Error (RMSE), Mean Absolute Error (MAE), Pearson Correlation Coefficient (PCC), and cosine similarity to assess the disparities between the raw count matrix, imputed data, and true count matrix. RMSE and MAE serve as indicators for measuring the numerical discrepancies between the imputed data and true expression data. A smaller value of RMSE and MAE indicates a more favorable performance of the imputation method. PCC and cosine similarity are numerical-based similarity measurement techniques used to evaluate the resemblance between imputed data and true expression data. The closer these two indicators approach 1, the more effective is deemed to be the imputation method employed in this study. It should be noted that due to dimension reduction in scGCC-imputed data compared with the true count matrix, calculation of corresponding indicators for this method is not feasible.

The results of the statistical indicators mentioned above, obtained from employing different imputation methods to process data with varying dropout rates, are presented in Figure 3. Specifically, scIDPMs(MHA) and scIDPMs(LSHA) represent the outcomes derived from networks constructed using multi-head attention and LSH attention respectively. As shown in the figure, scIDPMs consistently outperforms the other seven methods on simulated datasets with diverse dropout rates. Notably, as the dropout rate increases, other interpolation methods do not exhibit substantial improvement relative to the original data; however, scIDPMs maintains robustness and achieves optimal results throughout. It is worth mentioning that kNN-smoothing introduces more noise during interpolation by altering all expressed values; thus leading to larger corresponding RMSE and MAE values when fewer dropout positions occur (i.e., lower dropout rate). Furthermore, Figure 3 reveals no significant disparity in results between utilizing either of the two attention mechanisms.

**Figure 3.**
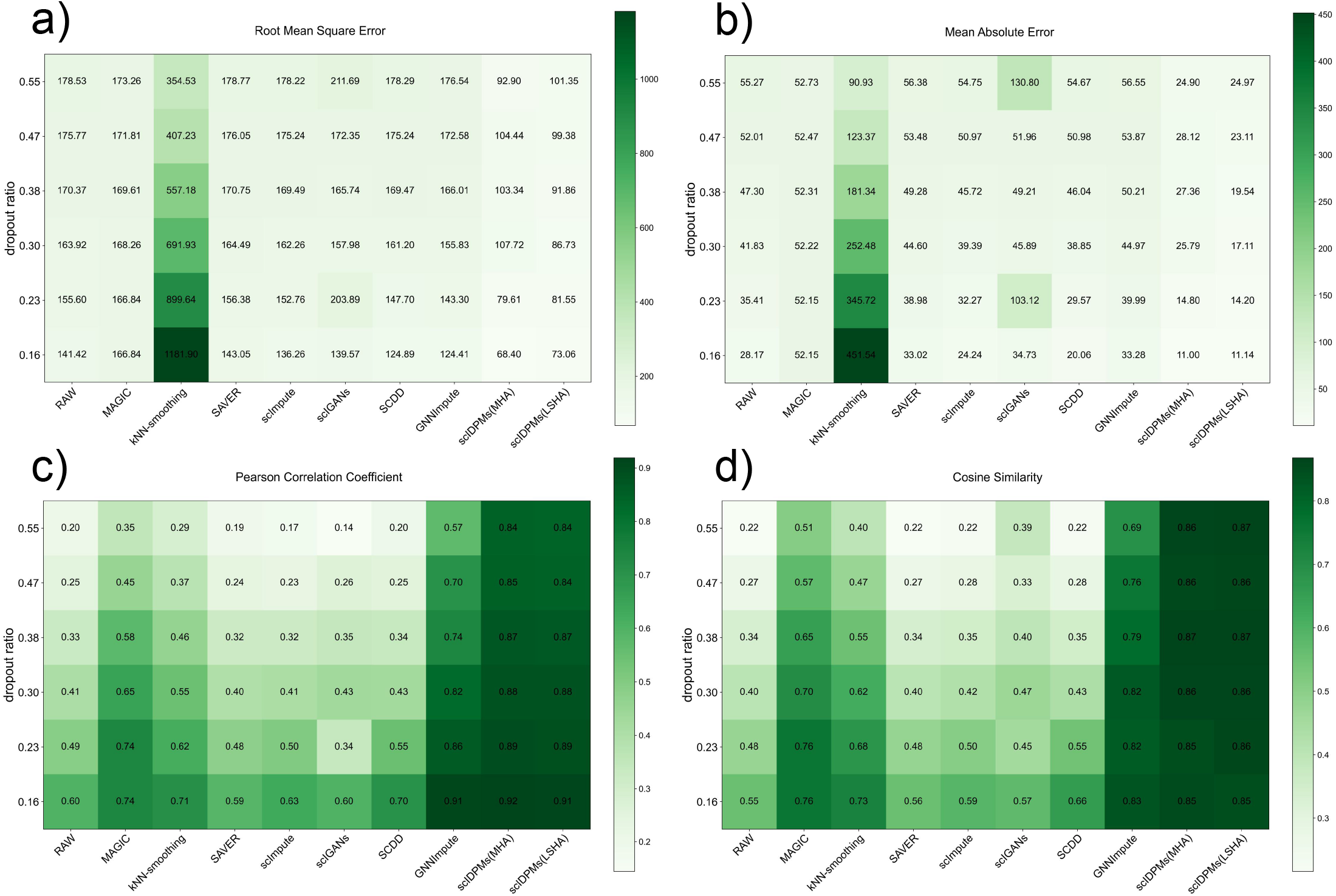
Performance of imputed data obtained by different methods to recover gene expression with different dropout rates.(**a-d**)RMSE MAE PCC cosine similarity of raw and imputed data with true counts matrix, respectively.

The UMAP visualization results of the raw data, true expression matrix, and imputed data with dropout rates of 0.16, 0.38, and 0.55 based on the real label of each cell are presented in Figure 4-6. Notably, scIDPMs exhibit distinct cell subgroups corresponding to the true count matrix at different dropout rates, reaffirming its reliability as a method for recovering missing gene expression values caused by dropout events without introducing supplementary noise. However, it is important to acknowledge that the introduction of additional noise during imputation by MAGIC and kNN-smoothing methods obscures genuine biological variation in the visualization outcomes of imputed data compared to those obtained from raw data. Although statistical indicators in Figure 3 do not show significant improvement when comparing scImpute and scIGANs with raw data, the imputation results for various dropout rates still demonstrate clearly distinguishable cell subgroups.

**Figure 4.**
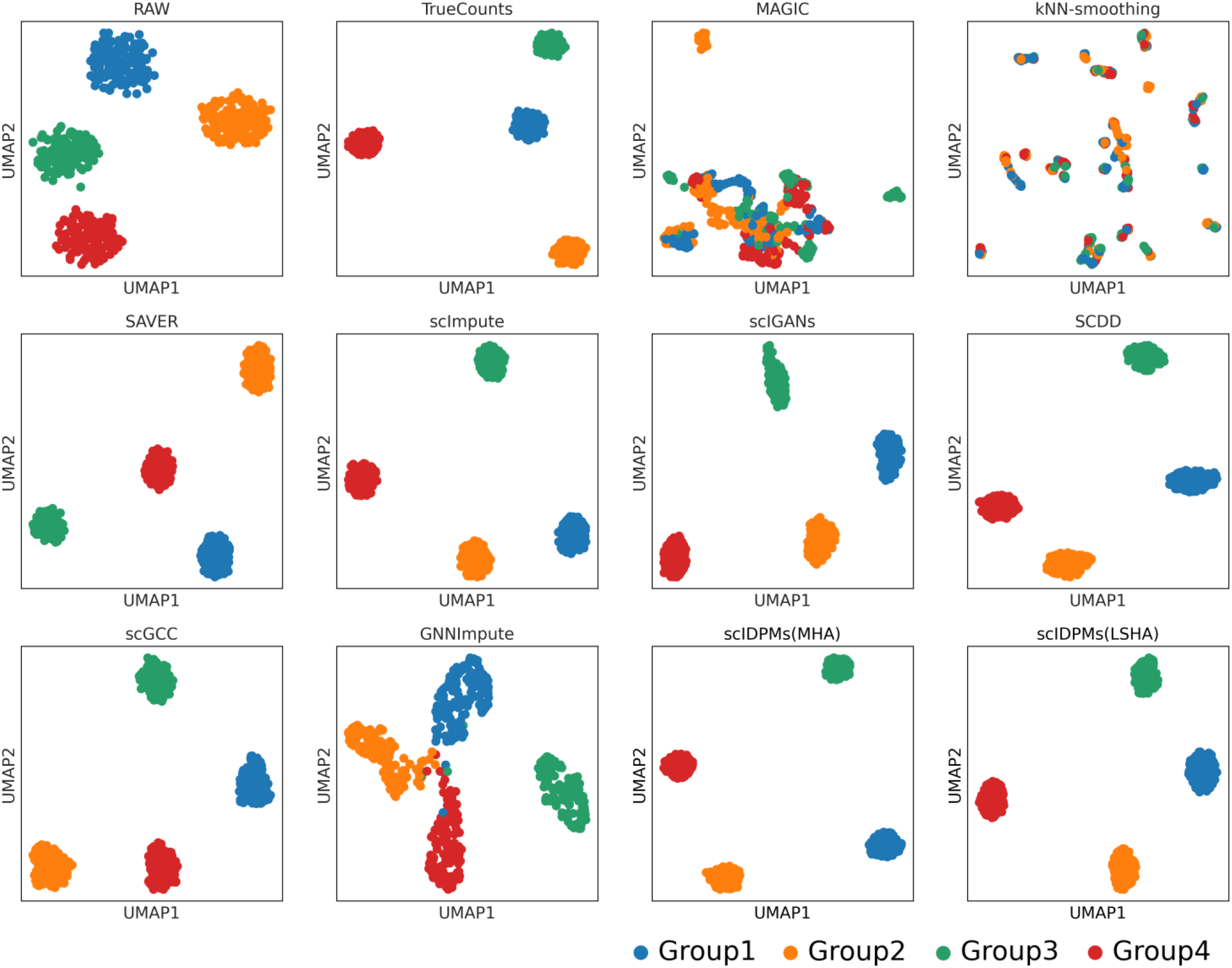
UMAP plots on raw data(dropout rate at 0.16), true counts matrix (without dropout), and imputed data by 10 methods.

**Figure 5.**
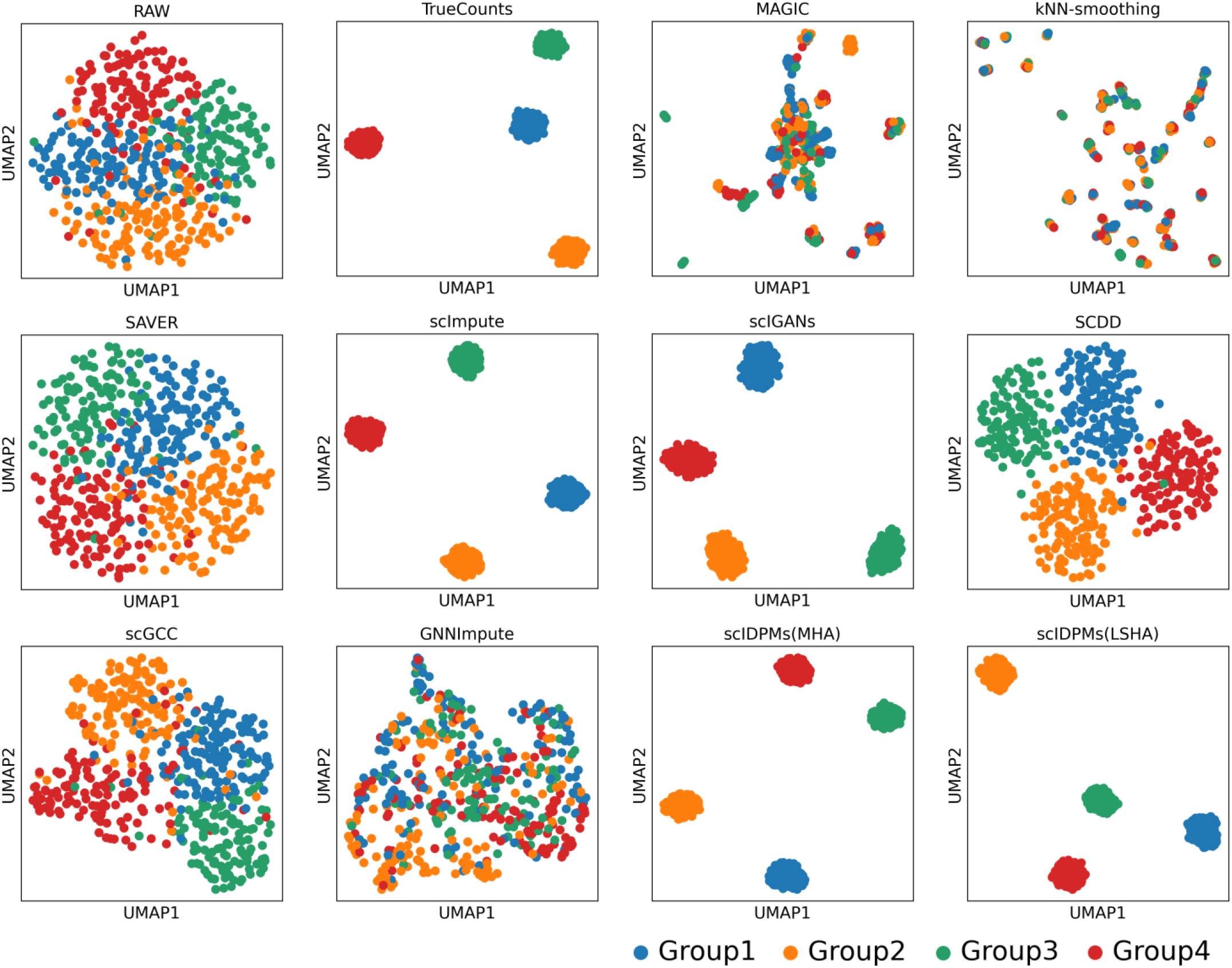
UMAP plots on raw data(dropout rate at 0.38), true counts matrix (without dropout), and imputed data by 10 methods.

**Figure 6.**
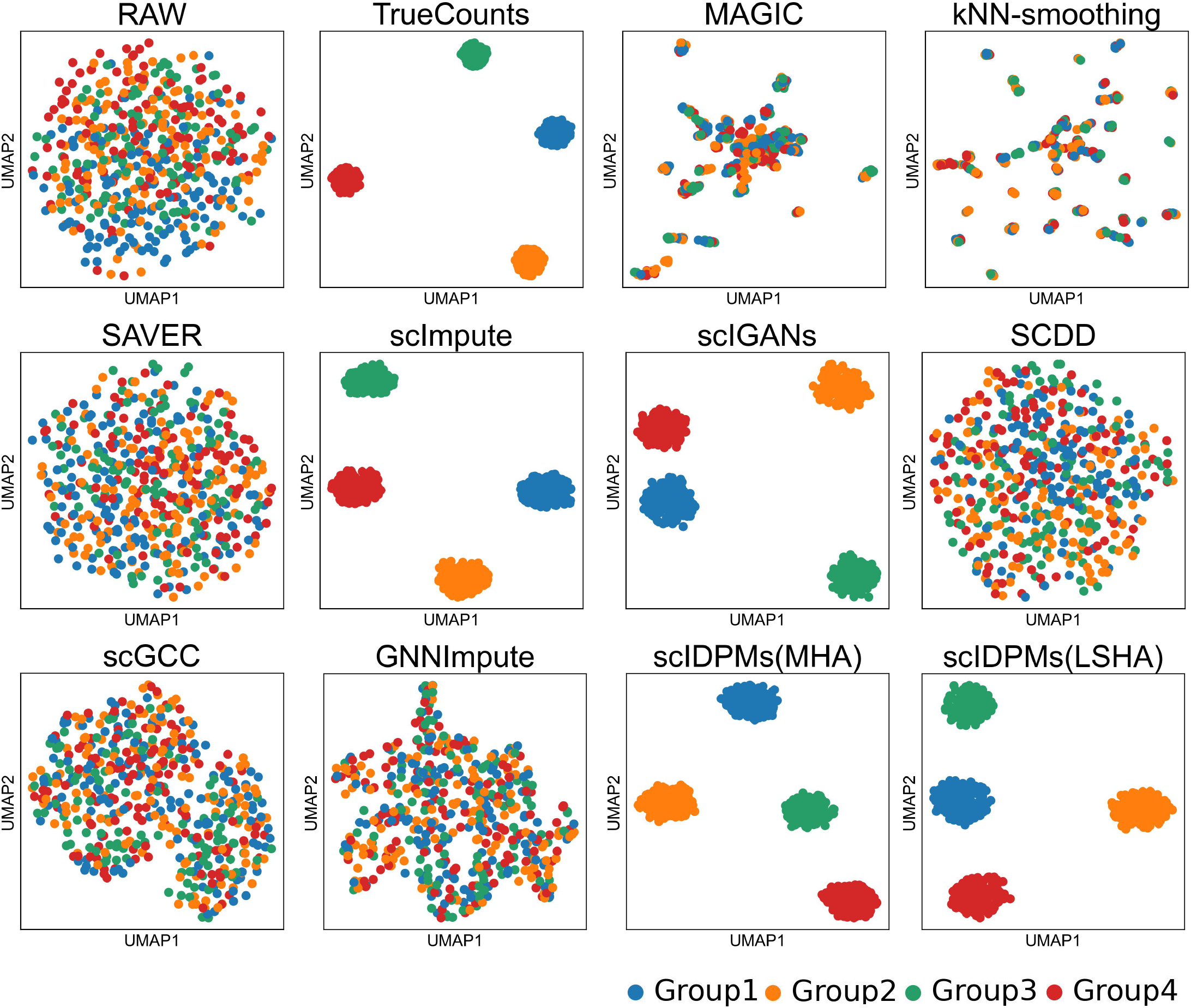
UMAP plots on raw data(dropout rate at 0.55), true counts matrix (without dropout),and imputed data by 10 methods.

### 3.2 scIDPMs facilitates the identification of distinct cell subpopulations

The process of cell clustering involves the utilization of clustering algorithms to categorize or group cells in scRNA-seq data based on their gene expression patterns or characteristics. The resulting clusters provide crucial insights for subsequent analyses, such as cell subgroup analysis and differential expression analysis, while also facilitating the discovery of novel cell types. An effective imputation method should possess the ability to accurately assign similar cells to the same cluster, thereby minimizing misclassification errors. To compare the performance of different methods in cell clustering, this paper utilizes the Mouse bone marrow mesenchyme (BM-MSCs) dataset published by Han et al. [46]. The count matrix derived from Gene Expression Omnibus (GEO) with access ID GSE108097 consists of 7365(cells) *×* 16322(genes) across 19 cell types, exhibiting high sparsity and dimensionality with a zero expression rate of 0.95. As each cell is labeled with its respective type, it provides an ideal resource for comparing the consistency and accuracy of clustering information and real class information after imputation.

To mitigate the impact of various factors and eliminate low-quality data, this study adopts the preprocessing and quality control method proposed by Zheng et al. [45] . Initially, genes with a count greater than 1 are retained, followed by identification of the top 1000 highly variable genes based on gene dispersion. The expression matrix is then filtered according to these calculations. This preprocessing approach not only reduces data noise but also preserves all single-cell samples while reducing sample dimensionality and addressing sequencing depth discrepancies. The preprocessed data serve as initial input for all imputation methods, which are subsequently used for clustering analysis. Specifically, we employ Scanpy’s built-in Leiden algorithm to cluster the interpolated data. Subsequently, differential expression analysis using the Wilcoxon rank-sum test identifies marker genes within each cluster, enabling labeling of cell subgroups in accordance with available dataset information. This evaluation allows us to assess the performance of the scIDPMs method in cell clustering and compare it against alternative methodologies.

The similarity between clustering results and real labels is assessed using four evaluation metrics, namely Adjusted Rand Index (ARI), F-score (also known as F1-score or F-measure), Purity, and Normalized Mutual Information (NMI). ARI provides a robust measure of similarity that takes into account the size of the dataset and the number of labels. F-score considers both accuracy and recall in classification outcomes. Purity primarily evaluates the homogeneity of clustering results. NMI incorporates both intracluster similarity and intercluster dissimilarity in assessing clustering outcomes. The evaluation metric results are presented in Table 2, where higher values indicate superior clustering performance. Unfortunately, scGCC failed to execute on this particular dataset. From Table 2, it can be observed that scIDPMs outperform other interpolation methods across various metrics, yielding the most favorable outcomes. With the exception of scImpute, no significant improvement in clustering results is observed with other imputation methods compared to raw data; some methods even exhibit poorer performance than raw data.

**Table 2.** Clustering result running on mouse bone marrow dataset.

|  | ARI | F-score | Purity | NMI |
| --- | --- | --- | --- | --- |
| RAW | 0.66 | 0.80 | 0.83 | 0.75 |
| MAGIC | 0.48 | 0.66 | 0.75 | 0.59 |
| kNN-smoothing | 0.46 | 0.47 | 0.59 | 0.56 |
| SAVER | 0.53 | 0.68 | 0.80 | 0.67 |
| scImpute | 0.81 | 0.90 | 0.91 | 0.85 |
| scIGANs | 0.66 | 0.78 | 0.83 | 0.75 |
| SCDD | 0.68 | 0.79 | 0.83 | 0.74 |
| GNNImpute | 0.43 | 0.61 | 0.74 | 0.62 |
| scIDPMs(MHA) | **0.98** | **0.99** | **0.99** | **0.98** |

The UMAP visualization results for both raw and imputed data based on real tags are presented in Figure 7. It is evident from the figure that even scImpute and scIGANs, which performed well on simulated data, were not satisfactory when applied to complex real scRNA-seq data. The imputed data failed to adequately restore gene expression characteristics, resulting in numerous cell samples with incorrect class information. In contrast, scIDPMs successfully clustered cell samples belonging to the same subgroup closely together, with only a few cells having misclassified labels. These visualized outcomes align with the findings reported in Table 2, indicating that scIDPMs effectively impute missing gene expression values while accurately preserving intercellular similarities and differences, thereby significantly enhancing the accuracy of subgroup clustering results.

**Figure 7.**
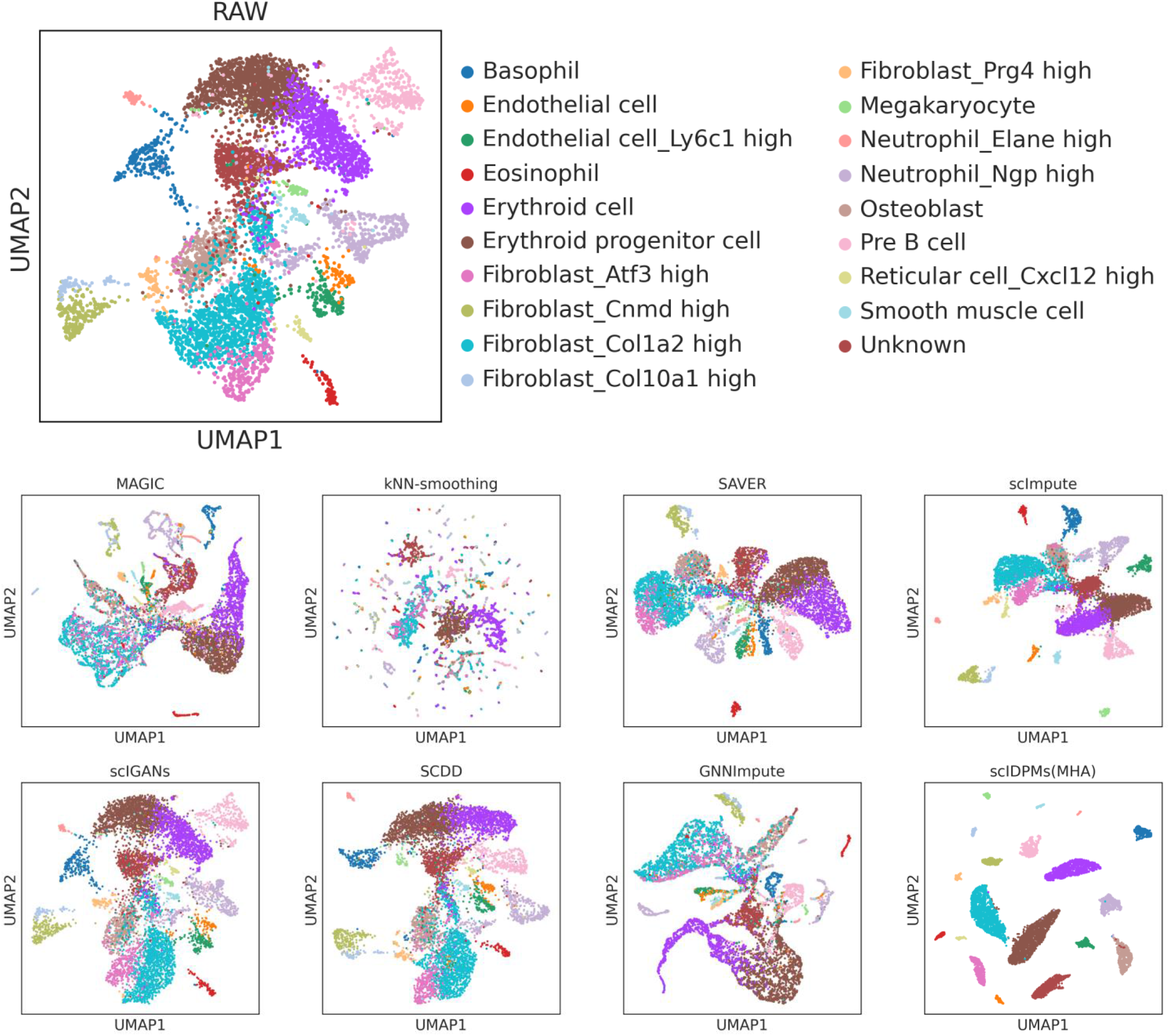
UMAP plots of raw and imputed data based on real cell categories on the mouse bone marrow dataset(GSE 108097).

### 3.3 scIDPMs improves the differential analysis

Differential analysis aims to identify genes exhibiting significant differential expression by comparing gene expression levels between different cell types or within the same cell type. This process plays a crucial role in identifying distinct cell types and studying mechanisms underlying gene expression regulation. In this study, we utilized the reduced 68kPBMCs dataset integrated into Scanpy to assess the efficacy of imputation methods in enhancing differential analysis performance. The dataset comprises 7 cell types (B-cell, Dendritic, Monocytes, NK, Other, Plasma, T-cell). Compared to the original PBMC 68k dataset, only 724 cells and 221 highly variable genes were retained along with authentic cell tags. This facilitates visualization of marker gene expression and analysis of genes displaying high variability. Marker genes refer to genes that are highly expressed in specific cell types and are commonly used by researchers for determining cell types. To evaluate the effectiveness of imputation methods on marker gene expression within each cell type, we compared the original data with data interpolated by scIDPMs as shown in Figure 8 a. Notably, after scIDPMs imputation, there is a more pronounced difference in marker gene expression between different cell types. Particularly noteworthy is that in the ‘Other’ type of cells, the expression level of IGLL1 gene significantly increases compared to the original data indicating enhanced specificity.

**Figure 8.**
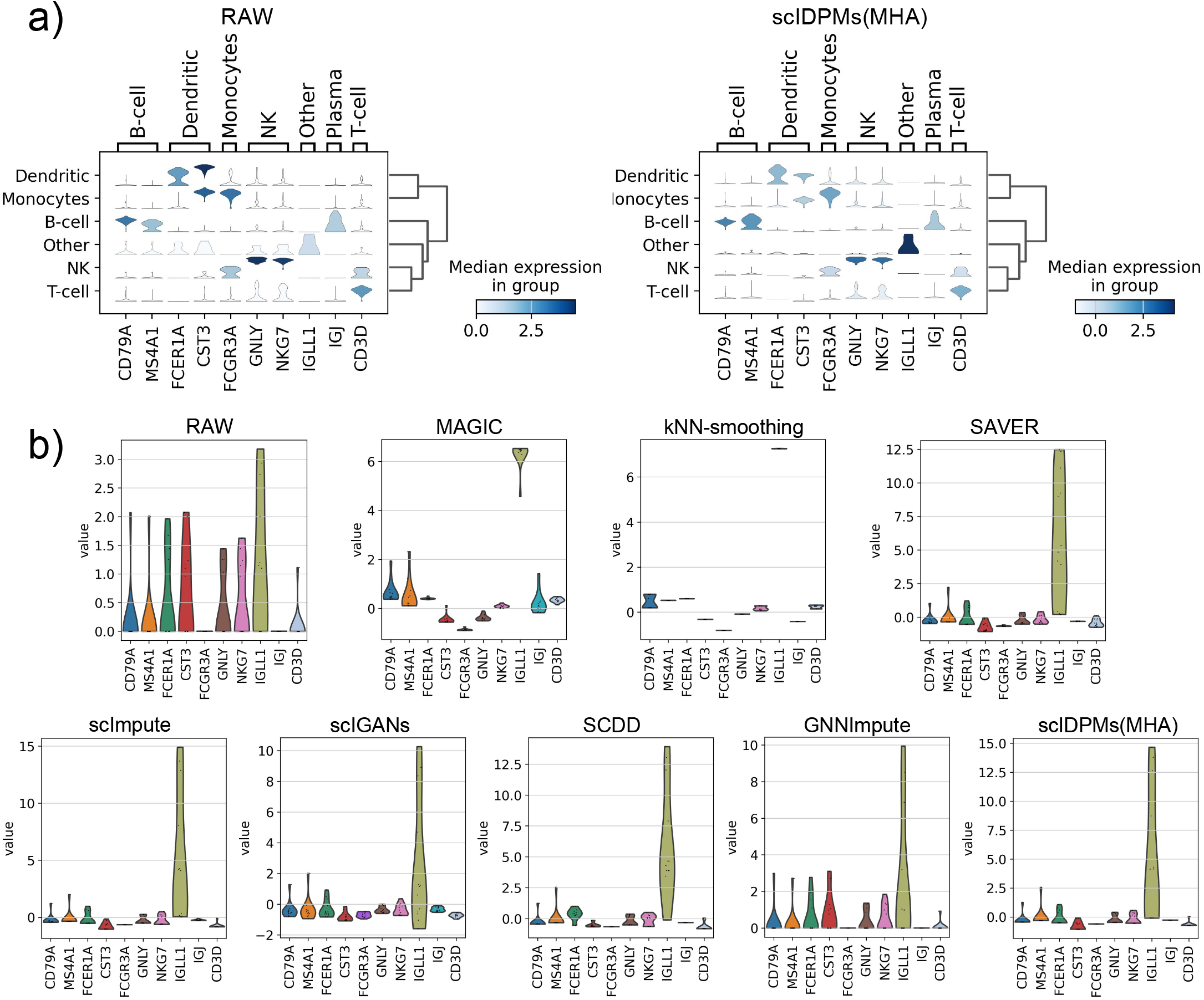
Visualization of differential analysis results between raw and imputed data in the reduced 68k PBMCs dataset.(**a**) Visualize the expression of marker genes in each cell type based on the raw data and imputed data by scIDPMs. (**b**)Visualization of expression distribution of marker genes in the ‘Other’ cell type based on raw data and imputed data by different methods.

To visually validate the effectiveness of scIDPMs method furtherly, we analyzed the distribution of marker gene expressions specifically within ‘Other’ type cells using various imputation methods (Figure 8 b). It should be noted that scGCC failed to run on this dataset due to technical limitations. The results demonstrated that both scImpute and scIDPMs outperformed other methods by effectively restoring IGLL1 gene deletion and enabling more accurate differentiation of ‘Other’ cell type during differential analysis.

The Wilcoxon rank-sum test method was employed to calculate differentially expressed genes within each cell type for both the raw data and the data after scIDPMs imputation. Figure 9 illustrates the top 4 differentially expressed genes in each cell type. It is evident that the scIDPMs method effectively fills in missing expression values across various cell types, thereby enhancing the significance of differential expression among highly expressed genes within each specific cell type compared to those outside it. Furthermore, when compared to the raw data, imputed data using scIDPMs enables detection of a greater number of highly expressed genes, such as TYROBP gene in ‘Monocytes’ cells and GZMB gene in ‘NK’ cells. These experimental findings demonstrate that scIDPMs can enhance accuracy and reliability in differential analysis, providing researchers with more precise insights into single-cell gene expression differences.

**Figure 9.**
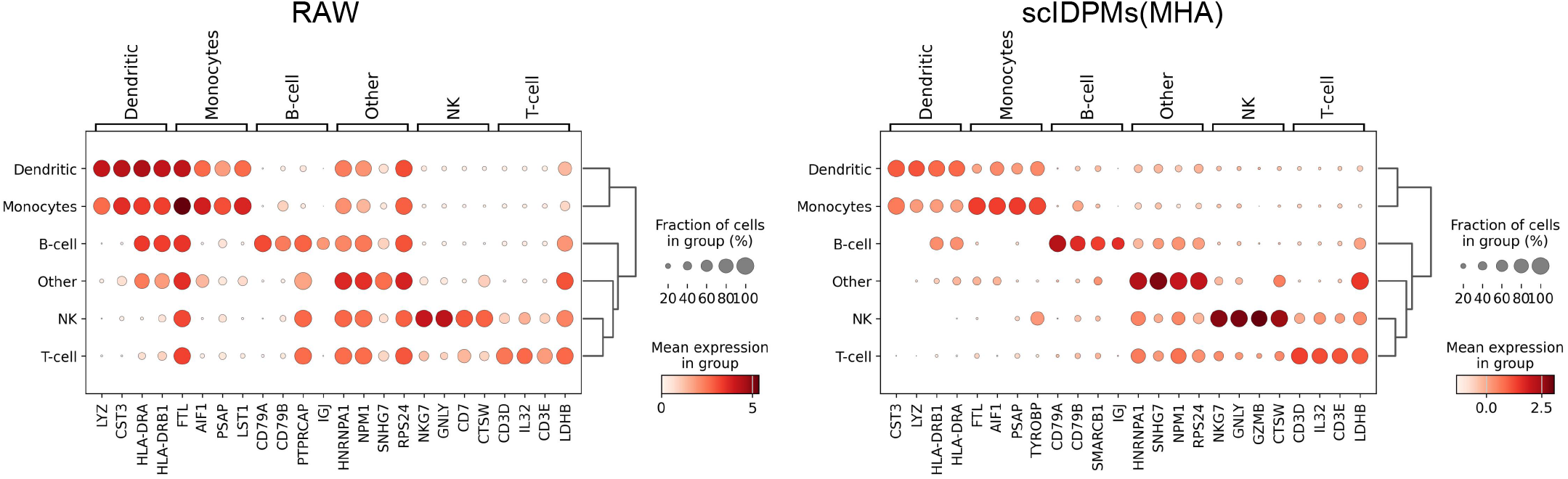
Visualization of the top 4 differentially expressed genes for each cell type based on the raw data and imputed data by scIDPMs.

## 4 CONCLUSION

To address the issue of missing gene expression values caused by dropout events in scRNA-seq data, this paper proposes a novel imputation method called scIDPMs, which is based on conditional diffusion probability models (DPMs). The proposed approach utilizes contextual information from the dropout site to infer the missing gene expression values. Initially, scIDPMs predicts the dropout site using gene expression data from cells belonging to the same class and then employs a self-supervised training approach to train conditional DPMs by dividing existing gene expression values into conditional observations and imputation targets. Subsequently, all available gene expression values are used as conditional observations to restore the ground-true distribution of dropout values from standard Gaussian noise using trained conditional DPMs. Compared with other imputation methods, scIDPMs achieves more accurate restoration of missing value distributions through a sampling algorithm based on DPMs. Furthermore, scIDPMs incorporates an attention mechanism that effectively captures global characteristics and key information within gene expression sequences.

In this study, simulated and real scRNA-seq data are utilized for evaluating the performance of scIDPMs in restoring gene expression values as well as improving cell clustering accuracy and differential analysis outcomes. Extensive experimental results demonstrate that compared to eight alternative imputation methods, scIDPMs consistently outperforms them across various datasets. Additionally, we explore different model architectures using multi-head attention mechanism and locality sensitive hashing(LSH) attention mechanism respectively for constructing network models and compare their performances on imputation results; these findings provide valuable insights for future research on processing high-dimensional sequencing data.

Based on the findings of this study, there are several intriguing avenues for future research that warrant exploration. Considering the inherent stochasticity in gene expression observed in scRNA-seq, it is plausible that the method employed in this paper to annotate dropout sites may not comprehensively capture all instances of dropout values. Therefore, alternative approaches should be considered to annotate dropout sites and compare imputation results more comprehensively. Although DPMs effectively restore data distribution, their sampling speed is comparatively slower when compared to other generative models. Hence, it would be advantageous to explore various ODE solvers or optimization algorithms to expedite sampling and potentially leverage pre-trained models.

## References

[1] Kevin Stevenson and Vladimir N Uversky. Single-cell rna-seq: a next generation sequencing tool for a high-resolution view of the individual cell. Journal of Biomolecular Structure and Dynamics, 38(12):3730–3735, 2020.

[2] Saiful Islam, Amit Zeisel, Simon Joost, Gioele La Manno, Pawel Zajac, Maria Kasper, Peter Lönnerberg, and Sten Linnarsson. Quantitative single-cell rna-seq with unique molecular identifiers. Nature methods, 11(2):163–166, 2014.

[3] Cheng Jia, Yu Hu, Derek Kelly, Junhyong Kim, Mingyao Li, and Nancy R Zhang. Accounting for technical noise in differential expression analysis of single-cell rna sequencing data. Nucleic acids research, 45(19):10978–10988, 2017.

[4] Tallulah S Andrews and Martin Hemberg. Identifying cell populations with scrnaseq. Molecular aspects of medicine, 59:114–122, 2018.

[5] Florian Wagner, Yun Yan, and Itai Yanai. K-nearest neighbor smoothing for high-throughput single-cell RNA-Seq data.

[6] David Van Dijk, Roshan Sharma, Juozas Nainys, Kristina Yim, Pooja Kathail, Ambrose J. Carr, Cassandra Burdziak, Kevin R. Moon, Christine L. Chaffer, Diwakar Pattabiraman, Brian Bierie, Linas Mazutis, Guy Wolf, Smita Krishnaswamy, and Dana Pe’er. Recovering Gene Interactions from Single-Cell Data Using Data Diffusion. Cell, 174(3):716–729.e27, July 2018.

[7] Mo Huang, Jingshu Wang, Eduardo Torre, Hannah Dueck, Sydney Shaffer, Roberto Bonasio, John I. Murray, Arjun Raj, Mingyao Li, and Nancy R. Zhang. SAVER: Gene expression recovery for single-cell RNA sequencing. Nature Methods, 15(7):539–542, July 2018.

[8] Wei Vivian Li and Jingyi Jessica Li. An accurate and robust imputation method scImpute for single-cell RNA-seq data. Nature Communications, 9(1):997, March 2018.

[9] Daniel J Kiviet, Philippe Nghe, Noreen Walker, Sarah Boulineau, Vanda Sunderlikova, and Sander J Tans. Stochasticity of metabolism and growth at the single-cell level. Nature, 514(7522):376–379, 2014.

[10] Keiron O’Shea and Ryan Nash. An introduction to convolutional neural networks. arXiv preprint 1511.08458, 2015.

[11] Jie Zhou, Ganqu Cui, Shengding Hu, Zhengyan Zhang, Cheng Yang, Zhiyuan Liu, Lifeng Wang, Changcheng Li, and Maosong Sun. Graph neural networks: A review of methods and applications. AI open, 1:57–81, 2020.

[12] Umberto Michelucci. An introduction to autoencoders. arXiv preprint 2201.03898, 2022.

[13] Diederik P Kingma, Max Welling, et al. An introduction to variational autoencoders. Foundations and Trends® in Machine Learning, 12(4):307–392, 2019.

[14] Alankrita Aggarwal, Mamta Mittal, and Gopi Battineni. Generative adversarial network: An overview of theory and applications. International Journal of Information Management Data Insights, 1(1):100004, 2021.

[15] Divyanshu Talwar, Aanchal Mongia, Debarka Sengupta, and Angshul Majumdar. AutoImpute: Autoencoder based imputation of single-cell RNA-seq data. Scientific Reports, 8(1):16329, November 2018.

[16] Yungang Xu, Zhigang Zhang, Lei You, Jiajia Liu, Zhiwei Fan, and Xiaobo Zhou. scIGANs: Single-cell RNA-seq imputation using generative adversarial networks. Nucleic Acids Research, 48(15):e85–e85, September 2020.

[17] Chenyang Xu, Lei Cai, and Jingyang Gao. An efficient scrna-seq dropout imputation method using graph attention network. BMC bioinformatics, 22:1–18, 2021.

[18] Jian Liu, Yichen Pan, Zhihan Ruan, and Jun Guo. SCDD: A novel single-cell RNA-seq imputation method with diffusion and denoising. Briefings in Bioinformatics, 23(5):bbac398, September 2022.

[19] Snehalika Lall, Sumanta Ray, and Sanghamitra Bandyopadhyay. LSH-GAN enables in-silico generation of cells for small sample high dimensional scRNA-seq data. Communications Biology, 5(1):577, June 2022.

[20] Sheng-Wen Tian, Jian-Cheng Ni, Yu-Tian Wang, Chun-Hou Zheng, and Cun-Mei Ji. scgcc: Graph contrastive clustering with neighborhood augmentations for scrna-seq data analysis. IEEE Journal of Biomedical and Health Informatics, 2023.

[21] Zehao Xiong, Jiawei Luo, Wanwan Shi, Ying Liu, Zhongyuan Xu, and Bo Wang. scGCL: An imputation method for scRNA-seq data based on graph contrastive learning. Bioinformatics, 39(3):btad098, March 2023.

[22] Zimo Huang, Jun Wang, Xudong Lu, Azlan Mohd Zain, and Guoxian Yu. scGGAN: Single-cell RNA-seq imputation by graph-based generative adversarial network. Briefings in Bioinformatics, 24(2):bbad040, March 2023.

[23] Chichi Dai, Yi Jiang, Chenglin Yin, Ran Su, Xiangxiang Zeng, Quan Zou, Kenta Nakai, and Leyi Wei. scIMC: A platform for benchmarking comparison and visualization analysis of scRNA-seq data imputation methods. Nucleic Acids Research, 50(9):4877–4899, May 2022.

[24] Limin Xu, Jing Zhang, Yiqian He, Qianqian Yang, Tianhao Mu, Qiushi Guo, Yingqiang Li, Tian Tong, Shifu Chen, and Richard D Ye. Scrnapip: A systematic and dynamic pipeline for single-cell rna sequencing analysis. iMeta, page e132, 2023.

[25] Jascha Sohl-Dickstein, Eric A. Weiss, Niru Maheswaranathan, and Surya Ganguli. Deep Unsupervised Learning using Nonequilibrium Thermodynamics, November 2015.

[26] Jonathan Ho, Ajay Jain, and Pieter Abbeel. Denoising Diffusion Probabilistic Models, December 2020.

[27] Ling Yang, Zhilong Zhang, Yang Song, Shenda Hong, Runsheng Xu, Yue Zhao, Wentao Zhang, Bin Cui, and Ming-Hsuan Yang. Diffusion models: A comprehensive survey of methods and applications. ACM Computing Surveys, 2022.

[28] Cheng Lu, Yuhao Zhou, Fan Bao, Jianfei Chen, Chongxuan Li, and Jun Zhu. DPM-Solver: A Fast ODE Solver for Diffusion Probabilistic Model Sampling in Around 10 Steps, October 2022.

[29] Ashish Vaswani, Noam Shazeer, Niki Parmar, Jakob Uszkoreit, Llion Jones, Aidan N Gomez, Łukasz Kaiser, and Illia Polosukhin. Attention is all you need. Advances in neural information processing systems, 30, 2017.

[30] Tom Brown, Benjamin Mann, Nick Ryder, Melanie Subbiah, Jared D Kaplan, Prafulla Dhariwal, Arvind Neelakantan, Pranav Shyam, Girish Sastry, Amanda Askell, et al. Language models are few-shot learners. Advances in neural information processing systems, 33:1877–1901, 2020.

[31] Romal Thoppilan, Daniel De Freitas, Jamie Hall, Noam Shazeer, Apoorv Kulshreshtha, Heng-Tze Cheng, Alicia Jin, Taylor Bos, Leslie Baker, Yu Du, et al. Lamda: Language models for dialog applications. arXiv preprint 2201.08239, 2022.

[32] Yidong Ouyang, Liyan Xie, Chongxuan Li, and Guang Cheng. MissDiff: Training Diffusion Models on Tabular Data with Missing Values, July 2023.

[33] Etienne Becht, Leland McInnes, John Healy, Charles-Antoine Dutertre, Immanuel WH Kwok, Lai Guan Ng, Florent Ginhoux, and Evan W Newell. Dimensionality reduction for visualizing single-cell data using umap. Nature biotechnology, 37(1):38–44, 2019.

[34] Vincent A Traag, Ludo Waltman, and Nees Jan Van Eck. From louvain to leiden: guaranteeing well-connected communities. Scientific reports, 9(1):5233, 2019.

[35] Lilian Weng. What are diffusion models? lilianweng.github.io, Jul 2021.

[36] Yusuke Tashiro, Jiaming Song, Yang Song, and Stefano Ermon. CSDI: Conditional Score-based Diffusion Models for Probabilistic Time Series Imputation, October 2021.

[37] Kaiming He, Xiangyu Zhang, Shaoqing Ren, and Jian Sun. Deep residual learning for image recognition. In Proceedings of the IEEE conference on computer vision and pattern recognition, pages 770–778, 2016.

[38] Yoon Kim. Convolutional neural networks for sentence classification. arXiv preprint 1408.5882, 2014.

[39] Dzmitry Bahdanau, Kyunghyun Cho, and Yoshua Bengio. Neural machine translation by jointly learning to align and translate. arXiv preprint 1409.0473, 2014.

[40] Nikita Kitaev, Łukasz Kaiser, and Anselm Levskaya. Reformer: The efficient transformer. arXiv preprint 2001.04451, 2020.

[41] Adam Paszke, Sam Gross, Francisco Massa, Adam Lerer, James Bradbury, Gregory Chanan, Trevor Killeen, Zeming Lin, Natalia Gimelshein, Luca Antiga, et al. Pytorch: An imperative style, high-performance deep learning library. Advances in neural information processing systems, 32, 2019.

[42] F Alexander Wolf, Philipp Angerer, and Fabian J Theis. Scanpy: large-scale single-cell gene expression data analysis. Genome biology, 19:1–5, 2018.

[43] Alex Nichol and Prafulla Dhariwal. Improved Denoising Diffusion Probabilistic Models, February 2021.

[44] Wenpin Hou, Zhicheng Ji, Hongkai Ji, and Stephanie C. Hicks. A systematic evaluation of single-cell RNA-sequencing imputation methods. Genome Biology, 21(1):218, December 2020.

[45] Luke Zappia, Belinda Phipson, and Alicia Oshlack. Splatter: simulation of single-cell rna sequencing data. Genome biology, 18(1):174, 2017.

[46] Xiaoping Han, Renying Wang, Yincong Zhou, Lijiang Fei, Huiyu Sun, Shujing Lai, Assieh Saadatpour, Ziming Zhou, Haide Chen, Fang Ye, et al. Mapping the mouse cell atlas by microwell-seq. Cell, 172(5):1091–1107, 2018.

